# A genome-wide screen identifies genes that suppress the accumulation of spontaneous mutations in young and aged yeast cells

**DOI:** 10.1101/492587

**Authors:** Daniele Novarina, Georges E. Janssens, Koen Bokern, Tim Schut, Noor C. van Oerle, Hinke G. Kazemier, Liesbeth M. Veenhoff, Michael Chang

## Abstract

To ensure proper transmission of genetic information, cells need to preserve and faithfully replicate their genome, and failure to do so leads to genome instability, a hallmark of both cancer and aging. Defects in genes involved in guarding genome stability cause several human progeroid syndromes, and an age-dependent accumulation of mutations has been observed in different organisms, from yeast to mammals. However, it is unclear if the spontaneous mutation rate changes during aging, and if specific pathways are important for genome maintenance in old cells. We developed a high-throughput replica-pinning approach to screen for genes important to suppress the accumulation of spontaneous mutations during yeast replicative aging. We found 13 known mutation suppression genes, and 31 genes that had no previous link to spontaneous mutagenesis, and all acted independently of age. Importantly, we identified *PEX19*, encoding an evolutionarily conserved peroxisome biogenesis factor, as an age-specific mutation suppression gene. While wild-type and *pex19Δ* young cells have similar spontaneous mutation rates, aged cells lacking *PEX19* display an elevated mutation rate. This finding suggests that functional peroxisomes are important to preserve genome integrity specifically in old cells, possibly due to their role in reactive oxygen species metabolism.

**Author Summary:** Spontaneous mutations arise as a consequence of improper repair of DNA damage caused by intracellular (i.e. toxic by-products of normal cellular metabolism or inaccurate DNA replication) or external (e.g. UV light or chemotherapy) sources. Elevated mutagenesis is implicated in tumorigenesis, and an age-dependent accumulation of mutations has been observed in many organisms. However, it is still unclear how and at which rate mutations accumulate during aging. It is also unknown if specific mechanisms exist that protect the genome of aged cells. We developed a high-throughput, genome-wide approach to identify genes that suppress the accumulation of mutations during yeast replicative aging. Yeast replicative aging refers to the decline in viability a single cell experiences with increasing number of mitotic divisions. We identified a number of new genes that counteract the accumulation of mutations independently of age. Moreover, we discovered that *PEX19*, a gene involved in the biogenesis of peroxisomes, is important to prevent the accumulation of mutations in aged cells. Since *PEX19* is conserved in humans, our work might help understand how human cells could better protect their genome from mutations during aging.

## Introduction

Genomic instability, which refers to an increased rate of accumulation of mutations and other genomic alterations, is a hallmark, and a likely driving force, of tumorigenesis [1]. Genomic instability is also a hallmark of aging [2], as suggested by the age-related accumulation of mutations observed in yeast, flies, mice and humans [3], and highlighted by the fact that defects in DNA repair pathways result in human premature aging diseases [4]. However, whether genome instability has a causative role in aging is still controversial [3], and it is not known if aged cells rely more heavily on specific genome maintenance pathways.

*Saccharomyces cerevisiae* is a convenient model to study genomic instability and its relationship with aging, since genome maintenance pathways are evolutionary conserved [5,6] and a large number of genetic assays have been developed to study DNA repair and mutagenesis in budding yeast [6,7]. Furthermore, most aging-related cellular pathways, as well as lifespan-modulating environmental and genetic interventions, show a remarkable degree of conservation from yeast to mammals [8–10]. There are two main *S. cerevisiae* aging models: replicative aging refers to the decline in viability that a cell experiences with increasing number of mitotic divisions (a model for aging of mitotically active cells), while chronological aging refers to the decline in viability of a non-dividing cell as a function of time (a model for aging of post-mitotic cells) [11].

To identify genes and pathways involved in mutagenesis during yeast replicative aging, two main challenges need to be overcome. First, while *S. cerevisiae* is a leading model system for genetic and genomic studies and the availability of the yeast deletion collection makes this model organism particularly amenable for genome-wide genetic screens [12,13], cellular processes involving low-frequency events, such as point mutations, recombination events or gross chromosomal rearrangements, pose a specific technical challenge, since these events are barely detectable with standard genome-wide screening methods [14]. In a pioneering study, Huang and colleagues performed a genome-wide screen for yeast genes that suppress the accumulation of spontaneous mutations in young cells by screening patches of large numbers of cells on solid media [15]. Patches of each strain of the deletion collection were replica-plated on media containing canavanine to detect canavanine-resistant (Can^R^) colonies arising from spontaneous mutations at the *CAN1* locus. A similar approach was subsequently used in other screens for genes controlling genome integrity [5,16–18]. This strategy has proven to be effective but is extremely laborious. To overcome these limitations, we developed a screening strategy to detect low-frequency events, based on high-throughput replica pinning of high-density arrays of yeast colonies.

The second challenge to identify genes involved in mutagenesis during yeast replicative aging is the isolation aged cells, since they constitute a tiny fraction of an exponentially growing cell population. To allow the study of a cohort of aging mother cells, Lindstrom and Gottschling developed the Mother Enrichment Program (MEP), an inducible genetic system that prevents the proliferation of daughter cells (Fig 1A) [19]. Upon activation by estradiol, the Cre recombinase, which is under the control of a daughter-specific promoter, enters the nucleus and disrupts two genes essential for cell cycle progression (namely *UBC9* and *CDC20*), resulting in an irreversible arrest of daughter cells in G2/M, while mother cells are unaffected. Thus, in the absence of estradiol, MEP cells grow exponentially and form normal colonies on an agar plate, while upon addition of estradiol, linear growth occurs and microcolonies are formed. Occasionally, due to spontaneous mutations, the MEP is inactivated and cells become insensitive to estradiol: these cells are called “escapers” [19]. Escaper cells grow exponentially and form normal sized colonies even in the presence of estradiol.

**Fig 1.**
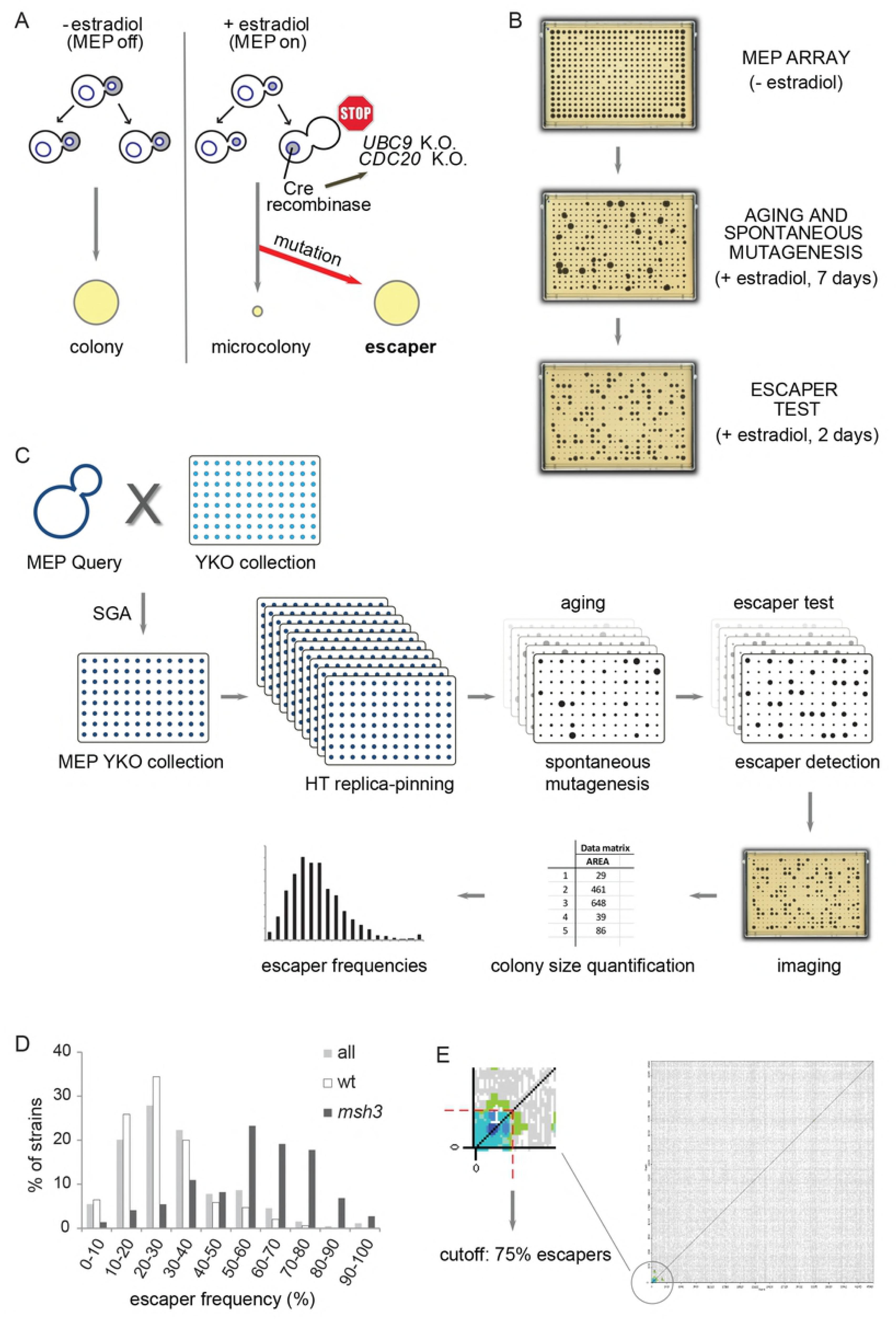
Combining the Mother Enrichment Program with high-throughput replica-pinning to screen for genes that suppress spontaneous mutations during yeast replicative aging. (A) The Mother Enrichment Program (MEP). Estradiol induction causes irreversible arrest of daughter cell proliferation, while growth of mother cells is unaffected. Inactivation of the MEP due to spontaneous mutations results in estradiol-insensitive cells called escapers. (B) Escaper formation is a readout for spontaneous mutation events during replicative aging. High-density arrays of MEP colonies are pinned on estradiol and escapers are subsequently detected by re-pinning on estradiol (big colonies). (C) Schematic of the screening procedure. The MEP is introduced into the YKO collection via Synthetic Genetic Array (SGA) methodology. The resulting MEP-YKO collection is amplified by high-throughput (HT) replica-pinning on estradiol, which activates the MEP and triggers replicative aging. A second replica-pinning on estradiol allows detection of escapers (escaper test). Escaper frequencies are calculated for each strain of the MEP-YKO collection. (D) Comparison of escaper frequencies of the whole MEP-YKO collection with escaper frequencies of wt and mutator (*msh3*) controls. (E) CLIK analysis sets an unbiased cutoff for validation. Green and blue colors indicate regions of the plot significantly enriched for physical and genetic interactions, while regions deprived of significant enrichment are plotted in gray. The red dotted line marks the cutoff suggested by the CLIK algorithm.

We combined the MEP system with a high-throughput replica-pinning strategy to perform a genome-wide screen aimed at identifying genes important to suppress the accumulation of spontaneous mutations in replicatively aging yeast cells, using escaper formation as a readout for spontaneous mutagenesis events. With our approach, we identified several new mutation suppression genes that act independently of age. We also found that *PEX19*, involved in peroxisome biogenesis, is an age-specific mutation suppression gene: while wild-type and *pex19Δ* young cells have similar spontaneous mutation rates, the absence of *PEX19* causes an elevated mutation rate specifically in old cells. We suggest that functional peroxisomes protect the genome of aged cells from spontaneous mutagenesis.

## Results

### A high-throughput screen to identify genes important for suppressing spontaneous mutations during yeast replicative aging

We developed a high-throughput replica-pinning strategy that enables detection of low-frequency events and used it to perform a genome-wide screen for genes important for suppressing spontaneous mutations during yeast replicative aging (Fig 1). We introduced the MEP system into the yeast knockout (YKO) collection via Synthetic Genetic Array (SGA) technology [20]. The resulting MEP-YKO collection was pinned multiple times in parallel on estradiol-containing plates (18 replicates per knockout strain) to activate the MEP, and grown for one week (Fig 1C). An example of our experimental setting is shown in Fig 1B. If at any time during aging a MEP-inactivating mutation occurs, an escaper colony is formed. Each plate was then re-pinned on estradiol and grown for two days to detect escapers (escaper test). At this point, all MEP-proficient mother cells that have exhausted their replicative potential are not able to give rise to a colony; conversely, if escaper cells are present, a fully grown colony can be observed (Fig 1B and C). Our high-throughput replica-pinning method allows the semi-quantitative estimation of spontaneous mutation rates on the basis of escaper formation (Fig 1C). Since every plate of the MEP-YKO collection was pinned multiple times in parallel on estradiol, it is possible to calculate the frequency of escaper formation for each deletion mutant strain in the collection: an increased escaper frequency compared to wild-type control strains is an indication of a high spontaneous mutation rate (Fig 1D).

To validate our assumption that the escaper frequency of each strain is a proxy for the spontaneous mutation rate, we could make use of the fact that 72 strains from the YKO collection were derived from a parental strain carrying an additional mutation in the mismatch repair gene *MSH3* and are therefore expected to show increased spontaneous mutation rates, independently of the identity of the knockout gene [21]. In addition, 340 empty positions randomly dispersed over the 14 plates of the MEP-YKO library were manually filled in with a wild-type MEP control. In Fig 1D, an overview of the escaper frequencies of the whole MEP-YKO collection is shown. Most of the strains have an escaper frequency between 10% and 40% (median: 27.8%). The wild-type control strains show a similar behavior (median: 22.2%), but with the important difference that the wild-type escaper frequency never exceeds 72.2%. In contrast, the escaper frequency of most of the *msh3* strains falls between 50% and 80% (median: 55.6%), validating the rationale of our screening method.

We then used Cutoff Linked to Interaction Knowledge (CLIK) analysis [22] to determine the cutoff for validation in an unbiased manner. The CLIK algorithm identified an enrichment of highly interacting genes at the top of our list (ranked according to escaper frequency), confirming the overall high quality of our screen (Fig 1E). The cutoff suggested by CLIK corresponds to an escaper frequency of 75%, which, not surprisingly, is slightly higher than the maximum escaper frequency observed in the wild-type controls (72.2%). To further explore the overall quality of the screen, we set the cutoff at 75% escaper frequency and performed phenotypic enrichment analysis using ScreenTroll, which examines the similarity between genome-scale screens [23]. Predictably, the first overlap with our gene list is the mutator screen performed by Huang and colleagues [15]. Furthermore, most of the top overlapping screens are related to genome instability and DNA damage sensitivity (S1 Table). Based on the threshold determined by CLIK, we proceeded to direct validation of all hits with an escaper frequency higher than 75%.

### Identification of new genes that suppress the accumulation of mutations independently of replicative age

We first discarded as false positive all hits where the escaper-causing mutation(s) had occurred before the beginning of the aging experiment (i.e. during the generation of the MEP-YKO library and before the subsequent high-throughput replica-pinning step) as in these cases, an escaper frequency of 100% is not an indication of an extremely high mutation rate. By spotting serial dilutions of strains from the MEP-YKO library on estradiol-containing plates (S1 Fig), we found that 25/115 hits had escaped before the actual screen started (S1 File). We then set out to validate the remaining 90 putative mutator strains.

Spontaneous mutations can occur at any moment of the replicative lifespan and our experimental design does not allow us to discriminate if a high escaper frequency is an indication of an increased mutation rate already in young cells, or of an elevated age-dependent accumulation of mutations. To distinguish between these two possibilities, we performed fluctuation tests to measure the forward mutation rate at the endogenous *CAN1* locus, where any type of mutation that inactivates the *CAN1* gene confers canavanine resistance [24,25]. Since fluctuation tests are performed with logarithmically growing cultures, they measure the spontaneous mutation rate in an age-independent fashion (i.e. in young cells). Twelve of our hits had been previously validated [15], and therefore were not re-tested. Several genes identified in the aforementioned study (namely *CSM2, SHU1, TSA1* and *SKN7*) fell just below our 75% escaper frequency cutoff (S1 File). Importantly, we validated 13 new mutator mutants. Our screening strategy thus enabled us to identify new genes important for the suppression of spontaneous mutations independently of replicative age. The 26 genes whose deletion results in an increased *CAN1* mutation rate of at least 1.8-fold compared to wild type are listed in Table 1. We named this group of genes “general mutation suppression genes” because the corresponding knockout strains, besides showing an elevated escaper frequency in our screening setup, also display an increased mutation rate when tested in young cells with a second assay for spontaneous mutagenesis at a different genetic locus. Of the general mutation suppression genes identified, 16/26 have one or more human orthologs. As expected, these genes are significantly enriched for Gene Ontology categories related to DNA damage response, DNA repair and recombination (Fig 2 and S2 File).

**Table 1.**
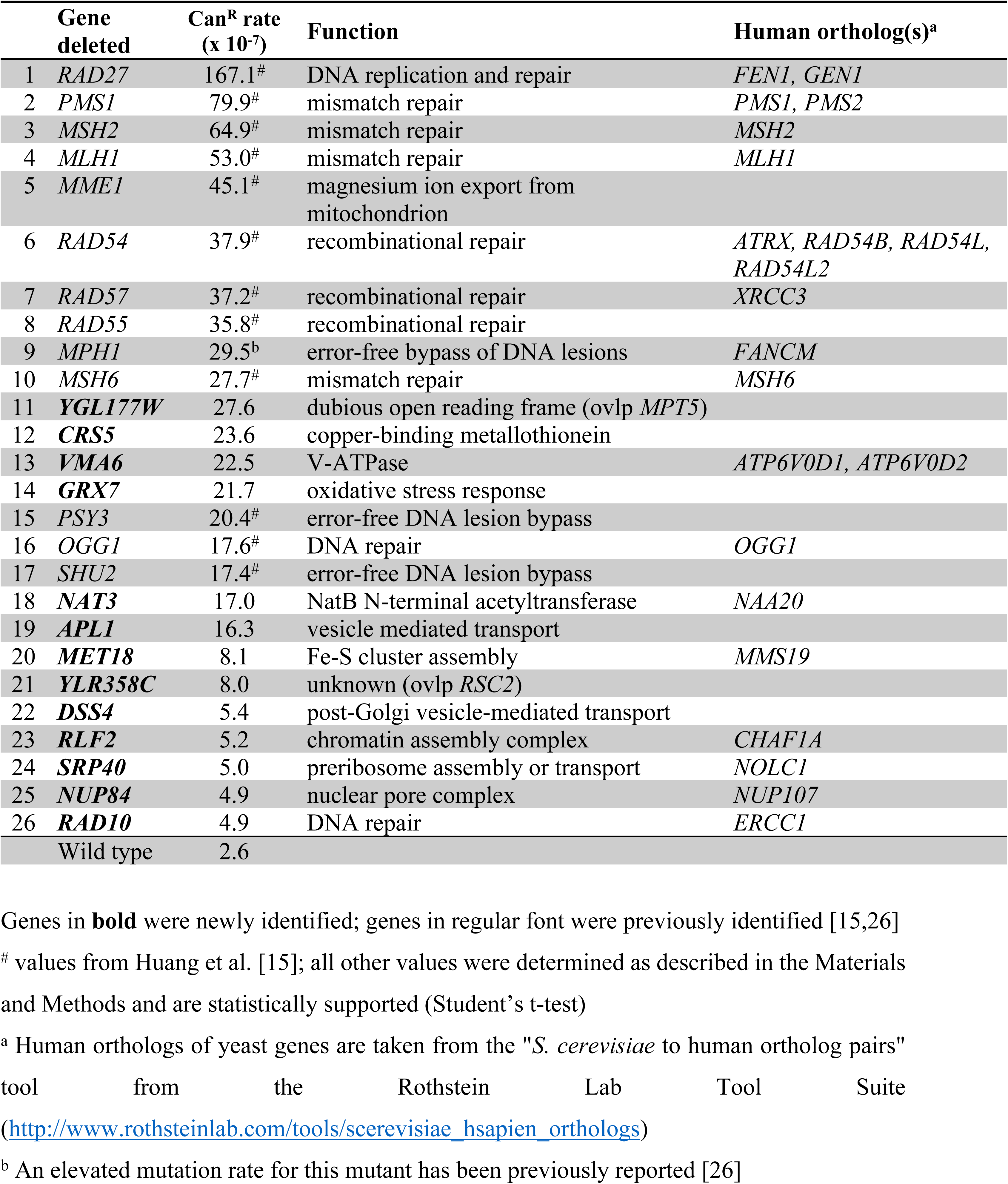
List of validated general mutation suppression genes.

**Fig 2.**
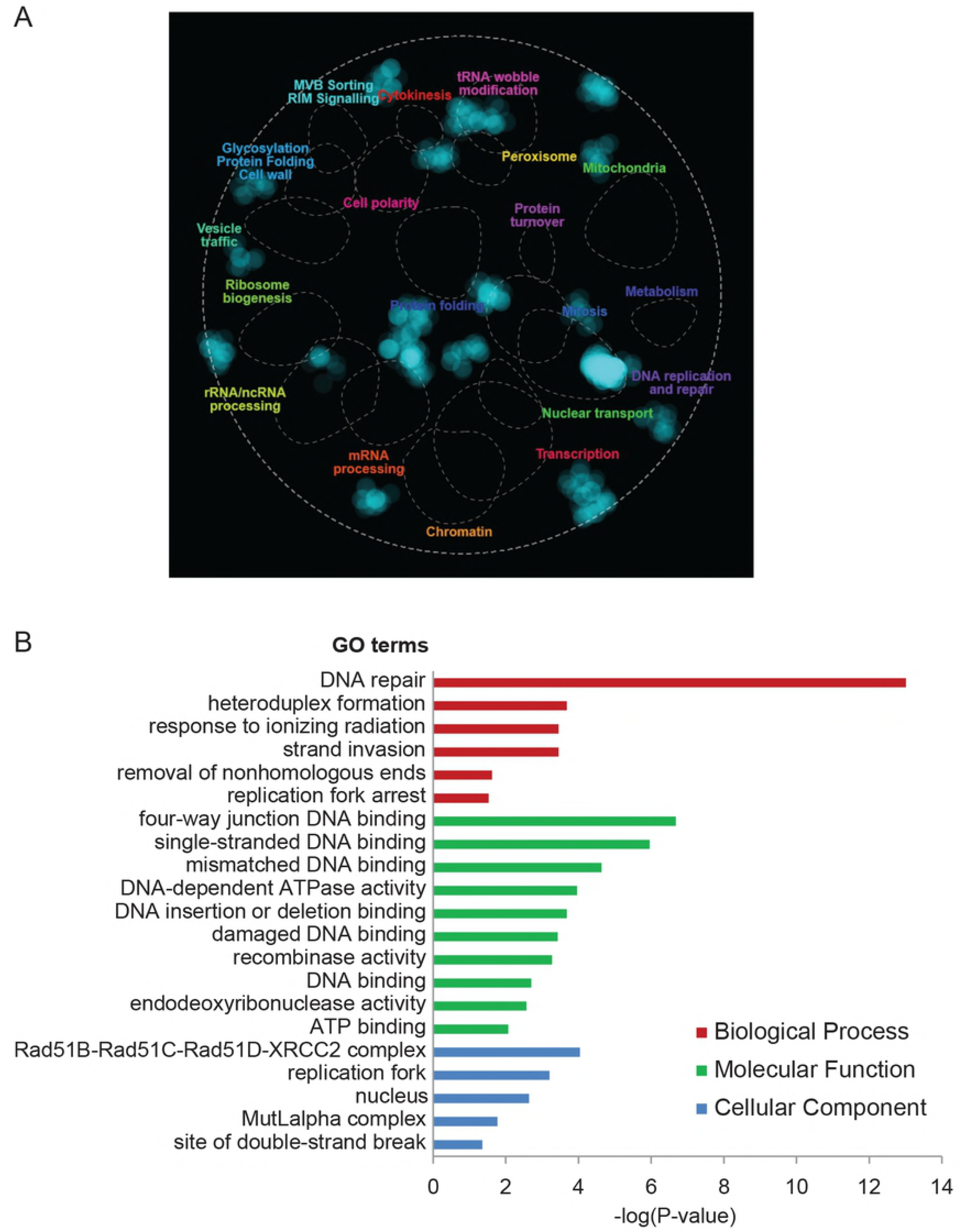
Overview of the general mutation suppression genes. (A) Functional enrichment in the yeast genetic landscape. Dotted lines indicate functional domains within the yeast genetic landscape, i.e. gene clusters enriched for a specific set of GO terms (the name of each functional domain is indicated by a colored label). Regions of the global similarity network significantly enriched for genes exhibiting genetic interactions with general mutation suppression genes were mapped using SAFE and are indicated in blue. (B) Gene Ontology enrichment analysis.

We then determined whether the 64 strains that do not show an elevated mutation rate at the *CAN1* locus would display an increase in the mutation rate if measured by escaper formation in the MEP genetic background. To do so, we performed a slightly modified version of the fluctuation test, where selective (i.e. estradiol-containing) plates are incubated for seven days and colonies are counted after two and seven days. Escaper colonies appearing after two days of incubation originate from mutations occurring prior to plating (i.e. in young cells), while all colonies appearing between day 2 and day 7 originate from a mutation event that occurred during replicative aging (see Materials and Methods for details). This experimental setup mimics the conditions in which the initial screen was performed and allows us to simultaneously measure the escaper formation rates in young cells and the age-dependent escaper formation frequencies. With this assay we identified 18 genes whose deletion results in an increased escaper formation rate of at least 1.8-fold compared to the wild type, independently of replicative age (i.e. based on colonies counted at day 2). We named these genes “MEP-specific mutation suppression genes”, since the spontaneous mutation rate measured at the *CAN1* locus in the corresponding knockout mutants is indistinguishable from the wild type (Table 2). About half (8/18) of these genes have one or more human orthologs. Intriguingly, MEP-specific mutation suppression genes are enriched for members of the THO/TREX complex, which is involved in co-transcriptional mRNA export from the nucleus. This process is important in the interplay between transcriptional elongation and R-loop formation in yeast and mammalian cells [27,28] (Fig 3 and S2 File).

**Table 2.**
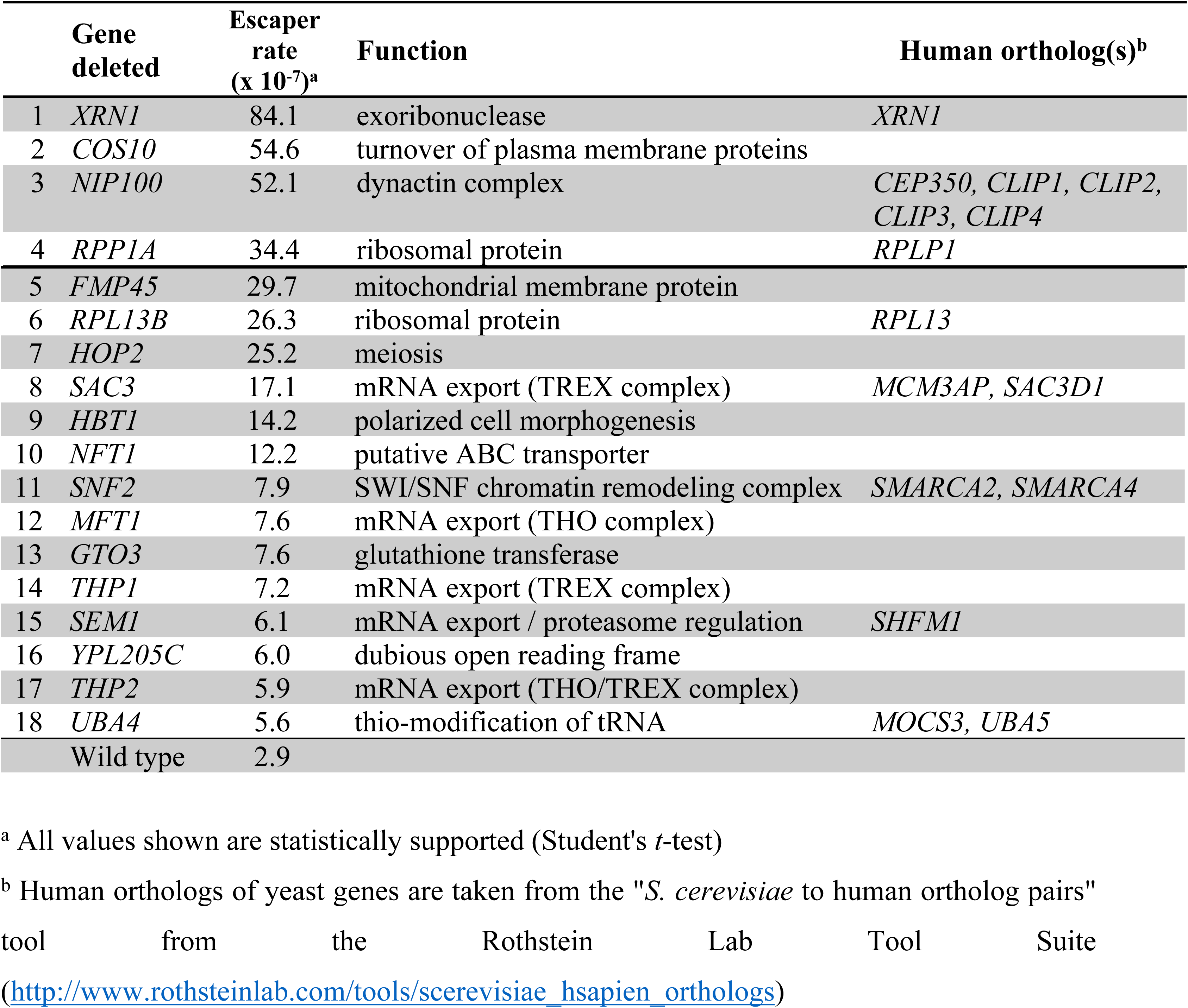
List of validated MEP-specific mutation suppression genes.

**Fig 3.**
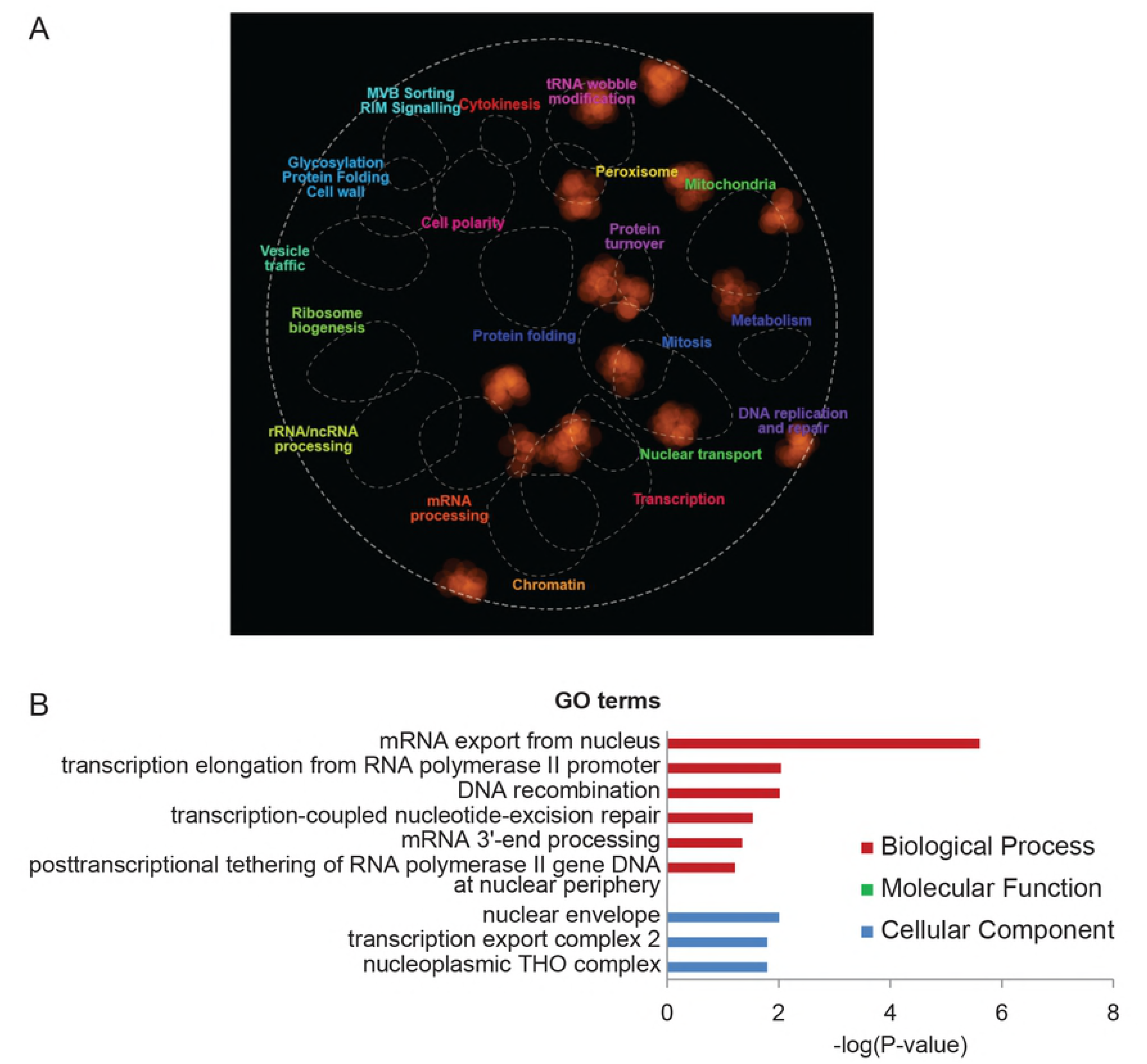
Overview of the MEP-specific mutation suppression genes. (A) Functional enrichment in the yeast genetic landscape. Regions of the global similarity network significantly enriched for genes exhibiting genetic interactions with MEP-specific mutation suppression genes were mapped using SAFE and are indicated in orange. See Legend to Fig 2A for details. (B) Gene Ontology enrichment analysis.

### *PEX19* suppresses age-dependent accumulation of mutations

At the end of our validation pipeline, we were left with four gene knockout strains that display no significant increase in forward mutation rate at the *CAN1* locus and in escaper formation rate in young cells (colonies counted at day 2) but show a higher age-dependent escaper frequency compared to wild type (colonies counted at day 7) (S2 Table). We were particularly interested in these genes, since our observations might indicate an age-dependent mutator phenotype. To validate these putative age-specific mutator mutants with an independent and more accurate method, we mechanically isolated young and aged mother cells by biotinylation and magnetic sorting and measured mutation frequencies at the *CAN1* locus in both cell populations [29]. Based on the Can^R^ frequencies in young cells, the replicative age (assessed by bud scar counting), and the mutation rate in young cells (previously determined by fluctuation test), we could calculate the expected Can^R^ frequencies in aged cells under the assumption that the mutation rate remains constant during replicative aging (see Materials and Methods for details). By comparing the observed and the expected frequencies, it becomes clear if the mutation rate of a given strain is constant or varies as cells age.

To establish a reference, we tested wild type cells. Strikingly, the observed mutation frequency in aged cells (replicative age ∼17) was lower than expected (Fig 4B and S2 Fig), suggesting a decrease in the spontaneous mutation rate during replicative aging. We then measured mutation frequencies in young and old cells from the four putative age-specific mutator strains. After the first test, three of these strains did not show any increase in age-dependent mutation frequency compared to the wild type and were therefore discarded as false positives (S3 Fig and S2 Fig). Conversely, age-dependent mutation frequency in the absence of *PEX19* was higher than in the wild type. We therefore repeated the test and confirmed that *pex19Δ* aged cells (replicative age ∼15.5) display much higher mutation frequencies than expected (Fig 4 and S2 Fig). This result suggests that *PEX19*, encoding an evolutionarily conserved factor required for peroxisome biogenesis [30], suppresses age-dependent accumulation of mutations. We observed a similar effect after the deletion of *PEX3*, which causes the same peroxisome biogenesis defect as observed in the absence of *PEX19*, namely lack of detectable peroxisomal structures [31]. Mutation frequency in aged *pex3Δ* cells (replicative age ∼13) was much higher than expected (S4 Fig), supporting the notion that functional peroxisomes contribute to genome maintenance during yeast replicative aging.

**Fig 4.**
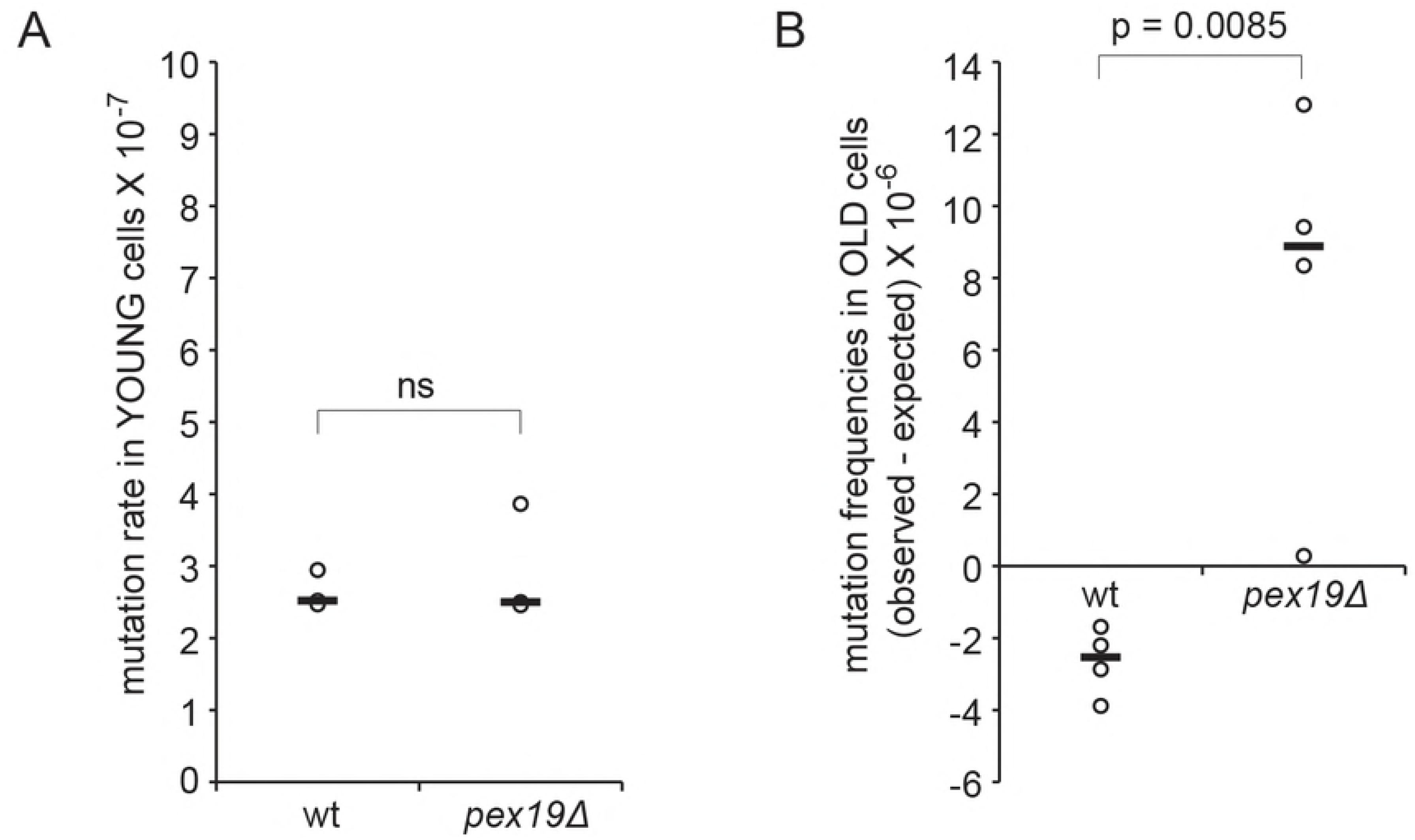
*PEX19* suppresses age-dependent accumulation of mutations. (A) *CAN1* forward mutation rate in young wt and *pex19Δ* cells. Values from three independent experiments are plotted. The thick dark bars represent the median values. ns: non-significant. (B) Age-dependent mutation frequencies at the *CAN1* locus in wt (replicative age ∼17) and *pex19Δ* (replicative age ∼15.5) cells. The difference between observed and expected mutation frequencies from four independent experiments is plotted. The horizontal bars represent the median values. p-value was determined by Student’s *t*-test.

## Discussion

*S. cerevisiae* is an outstanding model in which to perform genetic screens. However, there has been a lack of genome-wide, high-throughput screening techniques to detect low-frequency events (such as point mutations, recombination events and gross chromosomal rearrangements). These technical limitations, in combination with the difficulty to isolate large populations of replicative old cells, have so far hindered the study of genome maintenance during replicative aging.

### Potential applications of the high-throughput replica-pinning methodology

We developed a high-throughput replica-pinning approach to screen for cellular processes involving low-frequency events, thus filling a technological gap in the yeast screening field. We applied this strategy to screen for genes controlling the accumulation of spontaneous mutations during yeast replicative aging, using the Mother Enrichment Program both as a tool to induce replicative aging and as a reporter for spontaneous mutation events (Fig 1). Key technical adjustments (see Material and Methods) were: a) the use of 1536 format pads to pin the MEP-YKO collection in 384 format on estradiol, so that a smaller number of cells (∼1.5 x 10^4^) was deposited on the plate, to prevent nutrient limitation during replicative aging; b) the MEP-YKO collection amplification by parallel high-throughput replica-pinning to analyze 18 colonies per strain; c) the one-week incubation time, which allowed accumulation of spontaneous mutations throughout replicative lifespan. In this way, we were able to monitor enough cell divisions to detect low-frequency mutation events. Furthermore, the analysis of 18 independent colonies allowed the use of escaper frequency as a proxy for the spontaneous mutation rate. Bioinformatic analysis and experimental confirmation indicated the high quality of the screen, thus validating our methodology.

It is worth noting that, even independently of the MEP and the replicative aging perspective, a similar high-throughput replica-pinning approach can be used to screen for genes involved in other genome stability-related processes. For instance, we recently applied this strategy to study spontaneous homologous recombination events (manuscript in preparation). Similarly, our replica-pinning strategy could be adapted to screen for genes controlling genome integrity in the chronological lifespan model [32,33]. More generally, this technique can be used to study any process involving low-frequency events for which genetically selectable reporters exist or can be developed. Examples include transient (loss of) gene silencing [34], transcription errors [35] and read-through at premature termination codons [36].

### New general mutation suppression genes identified

Our screening setup was designed to allow simultaneous identification of age-independent and age-specific mutator mutants. We identified 13 new genes that suppress the accumulation of spontaneous mutations at the *CAN1* locus independently of age (Table 1). Some of these general mutation suppression genes (*RAD10* and *NUP84*) have defined roles in genome integrity [37–39]. For some other well-characterized genes, their role in preventing accumulation of mutations can be inferred from their molecular function. For instance, *MET18* has a conserved role in iron-sulfur (Fe/S) cluster assembly and insertion in several proteins involved in DNA replication and repair [40–42]. *RLF2*/*CAC1* encodes the largest subunit of the Chromatin Assembly Factor-I (CAF-1) complex, for which a role in DNA replication and repair of UV-induced DNA damage has been described [43–47].

For another group of new mutation suppression genes (*VMA6, GRX7, CRS5, NAT3*), their role in preventing accumulation of mutations might be more indirect. *VMA6* encodes a subunit of the evolutionary conserved vacuolar H^+^-ATPase (V-ATPase), responsible for vacuole acidification and cellular pH regulation [48,49]. Defects in yeast V-ATPase result in vacuole alkalinization and increased cytoplasm acidification [50], mitochondrial depolarization and fragmentation [51,52], altered iron homeostasis [53] and chronic endogenous oxidative stress [54]. This phenotype could potentially explain the elevated spontaneous mutation rate of a *vma6Δ* mutant, due to compromised functioning of Fe/S cluster-containing DNA replication and repair proteins as a consequence of mitochondrial depolarization [42,55], and/or to DNA damage caused by increased endogenous oxidative stress [56]. It would be interesting to test if disruption of other V-ATPase subunits causes the same mutator phenotype.

The *CRS5* gene product is a copper-and zinc-binding metallothionein [57,58]. The role of Crs5 in preventing spontaneous mutations is likely linked to its protective role against endogenous oxidative stress, since scavenging of reactive oxygen species is a general function of metallothioneins [59]. Crs5 might also directly protect DNA from copper-induced cleavage [60]. The reported physical interaction between Crs5 and the peroxiredoxins Tsa1 and Tsa2, responsible for preventing DNA damage and genome instability due to hydrogen peroxide and organic peroxides generated during normal cell metabolism, further supports a role for Crs5 in protecting the genome from oxidative damage [61,62].

Of particular interest is the mutation suppression genes *NAT3*, encoding the catalytic subunit of the conserved NatB N-terminal acetyltransferase. *NAT3* was identified in a screen for radiation sensitive mutants, and thereafter named *RAD56* [63,64]. Importantly, its human homologue hNAT3 has been implicated in carcinogenesis [65,66]. Non-degradative protein N-acetylation occurs co-translationally and can modulate protein folding, protein localization and protein-protein interactions [67]. It is likely that Nat3 prevents the accumulation of spontaneous mutations by ensuring the proper functioning of one (or more) of its targets. Interestingly, among the identified substrates of yeast NatB, several are involved in DNA metabolism, such as Pol31, Rnr4, Sml1, Nup84 [68–70]. Furthermore, many other factors involved in DNA processing and repair contain the peptide sequence recognized by NatB and are thus potential targets of Nat3 [71]. Further work will be needed to identify the relevant target(s) for preventing accumulation of spontaneous mutations.

*GRX7* encodes a largely uncharacterized glutaredoxin localized in the cis-Golgi [72,73]. How the disulfide bond-reducing activity of Grx7 in the Golgi affects spontaneous mutation rate in unclear, even though functional links between the Golgi apparatus and genome maintenance mechanisms have been suggested [74,75]. The *DSS4* gene product functions in the post-Golgi secretory pathway, while *APL1* is involved in clathrin-mediated vesicle transport. The elevated spontaneous mutation rate of *dss4Δ* and *apl1Δ* strains may indicate a thus far unanticipated connection between genome stability and vesicle transport in the secretory pathway. The remaining genes (*YGL177W, YLR358C*, and *SRP40*) are poorly characterized and require further investigation.

### MEP-specific age-independent mutation suppression genes are enriched for genes related to mRNA export

We also identified 18 mutants that, despite an undetectable increase in the spontaneous mutation rate at the *CAN1* locus, display an age-independent elevated escaper formation rate (Table 2). This observation hints at a locus-specific increase in mutagenesis for this group of mutator strains. The observation that MEP-specific mutation suppression genes are enriched for genes involved in mRNA export from the nucleus (Fig 3 and S2 File) suggests the involvement of R-loop-dependent genome instability [27,28]. R-loops form preferentially at specific genomic locations and can cause genomic instability by exposing single-stranded DNA tracts, triggering hyper-recombination and interfering with DNA replication [76]. Intriguingly, the exoribonuclease encoded by *XRN1*, our top MEP-specific mutation suppression gene, has also been implicated in preventing R-loop-dependent genome instability [77]. To confirm this hypothesis, one would need to examine the genomic features of the locus or loci where mutations that give rise to escapers happen. It is assumed that escaper-originating mutations occur at the *cre-EBD78* locus (since inactivating the Cre recombinase results in a disruption of the MEP system), but the creators of the MEP already suggested that this is not always the case, and other unknown endogenous loci might be involved in escaper formation [19]. Indeed, our genetic analysis of a few escapers originating from a wild-type MEP strain showed that the escaper phenotype does not always co-segregate with the *cre-EBD78* locus, confirming that mutations occurring at other genomic loci can result in MEP inactivation and escaper formation (S3 Table).

### A decrease in the spontaneous mutation rate in aged yeast cells

To investigate age-dependent spontaneous mutagenesis, we first compared mutation frequencies at the *CAN1* locus in young and old wt cells. Interestingly, *CAN1* mutation frequencies in aged cells are lower than predicted, indicating that spontaneous mutation rate decreases during replicative aging (Fig 4B). This might occur, for instance, if the efficiency of a mutagenic DNA repair pathway, such as translesion synthesis [78], is reduced in old cells. The same effect would be observed if an error-free repair pathway is upregulated during aging. A similar decrease in the spontaneous mutation rate at the *CAN1* endogenous locus has been previously reported, although the same study suggested that this effect could be locus-specific [29].

### A role for peroxisomes in suppressing age-dependent accumulation of spontaneous mutations

To better understand age-dependent mutagenesis, our screen aimed at identifying genes—if they exist—that prevent accumulation of spontaneous mutations specifically in old cells. We showed that *PEX19* is one of these genes, since its deletion has no effect on mutagenesis in young cells, but causes an elevated accumulation of mutations in aged cells (Fig 4). To our knowledge, this is the first described case of an age-dependent mutation suppression gene, suggesting that some cellular pathways are particularly important in protecting the genome of old cells.

*PEX19* is an evolutionary conserved gene which plays a key role in peroxisome biogenesis, and whose absence results in the lack of detectable peroxisomes [30,31]. Deletion of *PEX3*, another peroxisome biogenesis gene, causes the same age-specific mutator phenotype (S4 Fig), implying that functional peroxisomes are important to prevent age-dependent accumulation of mutations. The human orthologs of *PEX19* and *PEX3*, together with other peroxins, are mutated in Zellweger syndrome, a severe cerebro-hepato-renal peroxisome biogenesis disorder [79]. Peroxisomes are key organelles for the maintenance of the redox balance of the cell. On the one hand, they generate H_2_O_2_ as a consequence of fatty acid peroxidation; on the other hand, they contain a set of antioxidant enzymes and function therefore as reactive oxygen species (ROS) scavenging organelles [80]. Interestingly, loss of Pex19 in *D. melanogaster* causes ROS accumulation and mitochondrial damage, and a mouse model of Zellweger syndrome displays a similar phenotype [81,82]. Elevated endogenous ROS are known to induce genome instability [83,84].

How could peroxisomes potentially contribute to genome integrity maintenance in aged yeast cells? Several studies report an asymmetric ROS distribution between mother and daughter cell, resulting in ROS accumulation during replicative aging [85–87]. These elevated ROS levels are accompanied by an age-dependent hyperoxidation and inactivation of the peroxiredoxin Tsa1, a key antioxidant enzyme important for genome integrity and for suppression of mutations [15,62,83,84,88]. In this context, the role of peroxisomes in protecting the genome from endogenous oxidative stress might become crucial, due to the age-dependent increase in ROS accumulation and the concomitant progressive failure of other redundant antioxidant systems that are active in young cells. Our observations suggest that deletion of *PEX19* has a synergistic effect with age, resulting in elevated spontaneous mutagenesis. Given the evolutionary conservation of this and many other peroxisome biogenesis factors, it would be interesting to test the contribution of peroxisomes in genome maintenance during mammalian cell aging and cancer development. Indeed, several studies have reported the absence of peroxisomes in cancer cells [89–91], suggesting a possible link between peroxisome biogenesis defects and tumorigenesis [92].

## Materials and Methods

### Yeast strains and growth conditions

Standard yeast media and growth conditions were used [93,94]. All yeast strains used in this study are derivatives of the BY4741 genetic background [95] and are listed in Table 3. DNY34 was obtained from Y7092 and UCC8773 by crossing and tetrad dissection. The *ice2Δ::kanMX* strain from the deletion collection (EUROSCARF) strain was crossed with strain UCC8774 by standard yeast genetics to create the strain DNY80. Strains DNY99, DNY101, DNY102 and DNY105 were constructed by standard PCR-mediated gene deletion in strain UCC8773.

**Table 3.** Yeast strains used in this study.

| Strain name | Relevant genotype | Source |
| --- | --- | --- |
| BY4741 | *MATa his3Δ1 leu2Δ0 ura3Δ0 met15Δ0* | [95] |
| UCC8773 | *MATa his3Δ1 leu2Δ0 ura3Δ0 lys2Δ0 hoΔ::Pscw11-cre-EBD78-natMX loxP-CDC20-intron-loxP-hphMX loxP-UBC9-loxP-LEU2* | [96] |
| UCC8774 | *MATα his3Δ1 leu2Δ0 ura3Δ0 trp1Δ63 hoΔ::Pscw11-cre-EBD78-natMX loxP-CDC20-intron-loxP-hphMX loxP-UBC9-loxP-LEU2* | [96] |
| Y7092 | *MATα his3Δ1 leu2Δ0 ura3Δ0 can1Δ::STE2pr-Sp_his5 lyp1Δ met15Δ0* | [97] |
| DNY34 | *MATα his3Δ1 leu2Δ0 ura3Δ0 can1Δ::STE2pr-Sp_his5 lyp1Δ met15Δ0 hoΔ::PSCW11-cre-EBD78-natMX loxP-UBC9-loxP-LEU2 loxP-CDC20-Intron-loxP-hphMX* | This study |
| DNY80 | *MATa his3Δ1 leu2Δ0 ura3Δ0 lys2Δ0 met15Δ0 hoΔ::Pscw11-cre-EBD78-natMX loxP-CDC20-intron-loxP-hphMX loxP-UBC9-loxP-LEU2 ice2Δ::kanMX* | This study |
| DNY99 | *MATa his3Δ1 leu2Δ0 ura3Δ0 lys2Δ0 met15Δ0 hoΔ::Pscw11-cre-EBD78-natMX loxP-CDC20-intron-loxP-hphMX loxP-UBC9-loxP-LEU2 pex19Δ::kanMX* | This study |
| DNY101 | *MATa his3Δ1 leu2Δ0 ura3Δ0 lys2Δ0 met15Δ0 hoΔ::Pscw11-cre-EBD78-natMX loxP-CDC20-intron-loxP-hphMX loxP-UBC9-loxP-LEU2 rox3Δ::kanMX* | This study |
| DNY102 | *MATa his3Δ1 leu2Δ0 ura3Δ0 lys2Δ0 met15Δ0 hoΔ::Pscw11-cre-EBD78-natMX loxP-CDC20-intron-loxP-hphMX loxP-UBC9-loxP-LEU2 atg23Δ::kanMX* | This study |
| DNY105 | *MATa his3Δ1 leu2Δ0 ura3Δ0 lys2Δ0 met15Δ0 hoΔ::Pscw11-cre-EBD78-natMX loxP-CDC20-intron-loxP-hphMX loxP-UBC9-loxP-LEU2 pex3Δ::kanMX* | This study |

### High-throughput replica-pinning screen

High-throughput manipulation of high-density yeast arrays was performed with the RoToR-HDA pinning robot (Singer Instruments). The Mother Enrichment Program (MEP) was introduced into the *MAT***a** yeast deletion collection (EUROSCARF) through Synthetic Genetic Array (SGA) methodology [97] using the DNY34 query strain. The procedure was performed twice in parallel to generate two independent sets of MEP yeast deletion arrays in 384-colony format. When a specific MEP mutant was missing in one of the arrays, it was manually pinned over from the other set. Positions that were empty in both sets were filled with *his3*Δ*::kanMX* control strains, unless they were kept empty for plate identification purposes. Colonies from the two sets of MEP yeast deletion arrays were pinned onto YPD + G418 plates and incubated for six hours at 30°C. Each plate of each set was then pinned onto nine YPD plates containing 1 μM estradiol (18 replicates in total). At this step, colonies were pinned in 384 format using 1536 format pads, so that a smaller number of cells was deposited to prevent nutrient limitation.

Plates were incubated for seven days at 30°C and then scanned with a flatbed scanner. Subsequently, each plate was pinned onto one YPD plate containing 1 μM estradiol and incubated for two days at 30°C before scanning (“escaper test”). Colony area measurement was performed using the ImageJ software package [98] and the ImageJ plugin ScreenMill Colony Measurement Engine [99], to assess colony circularity and size in pixels. The data was filtered to exclude artifacts by requiring a colony circularity score greater than 0.8. Colonies with a pixel area greater than 200 were considered escapers, and for each deletion strain, the ratio of escapers to total colonies in replica pinning experiments was used as the escaper frequency score.

### Screen validation pipeline

Putative hits were initially analyzed by fluctuation test to measure the forward mutation rate at the endogenous *CAN1* locus. To do so, we used the strains from the YKO collection, because the strains from the MEP-YKO collection are *can1Δ*. At first, we performed one fluctuation test per strain. If the mutation rate was higher than 1.5-fold of the wild-type mutation rate, the test was repeated another two or three times.

For all the genes whose deletion does not cause an increase in the mutation rate at the *CAN1* locus, the corresponding knockout strains from the MEP-YKO collection were analyzed by fluctuation test to measure the escaper formation rate in young cells and the escaper formation frequency in replicatively aged cells. At first, we performed one fluctuation test per strain. If the escaper formation rate was higher than 1.5-fold of the wild-type escaper formation rate, the test was repeated another two or three times. In case no increase in escaper formation rate was detected but elevated age-dependent escaper formation frequencies were observed, the experiment was repeated another one or two times.

Strains that consistently displayed an elevated age-dependent escaper formation frequency were further validated by construction of a new knockout strain in a MEP *CAN1* background and by direct measurement of spontaneous mutation frequencies at the *CAN1* locus in young and aged cells. Each knockout strain was tested once. If the age-dependent mutation frequencies were not higher that the wild-type control, the strain was discarded as a false positive; if the age-dependent mutation frequencies were increased compared to the wild-type control, the experiment was repeated three times.

The identity of all validated strains from the YKO and MEP-YKO collections was confirmed by barcode sequencing as previously described [100].

### Measurements of the spontaneous forward mutation rate at the *CAN1* locus

Single colonies were inoculated in 5 ml YPD and grown up to saturation (two days at 30°C). 100 μl were plated onto canavanine-containing SD medium (50 μg/ml) to identify forward mutations in *CAN1* and 50 μl of a 10^5^-fold dilution was plated onto SD medium to count viable cells. Colonies were counted after two days of growth at 30°C and the spontaneous forward mutation rate at the *CAN1* locus was determined by fluctuation test from nine independent cultures using the method of the median [25,101]. Values represent the average of at least three independent experiments.

### Measurements of spontaneous escaper formation rate and age-dependent escaper formation frequencies

Single colonies were inoculated in 5 ml YPD and grown up to saturation (two days at 30°C). 50 μl of a 50-fold dilution (or a higher dilution, when needed) were plated onto YPD plates containing 1 μM estradiol to identify escaper occurrence in young cells (“young plates”), 50 μl of a 500-fold dilution (or a higher dilution, when needed) were plated onto YPD plates containing 1 μM estradiol to identify escapers occurrence in aging cells (“old plates”), and 50 μl of a 500000-fold dilution was plated onto YPD plates to count viable cell number. Colonies were counted after two days of growth at 30°C (for “young plates” and YPD plates) or after two and after seven days of growth at 30°C (for “old plates”). In “young plates”, colonies that were smaller than the colonies growing on the corresponding YPD plate were not counted, because for those colonies the escaper-causing mutation occurred after plating. The spontaneous escaper formation rate in young cells was determined by fluctuation test from 7-10 independent cultures using the MSS-maximum-likelihood estimator method from the FALCOR fluctuation analysis calculator [102]. Values represent the average of at least three independent experiments. Age-dependent escaper formation frequencies was calculated by dividing the number of escaper colonies that appeared between day 2 and day 7 (on “old plates”) by the number of viable cells plated (determined from the YPD plates).

### Measurements of spontaneous mutation frequencies at the *CAN1* locus in young and aged cells

Isolation of young and aged cells was performed essentially as previously described [19,103]. 1.5 x 10^9^ cells from a log-phase MEP culture were washed with cold phosphate buffered saline (PBS), resuspended in cold PBS containing 7 mg/ml Sulfo-NHS-LC-Biotin (Thermo Scientific) and incubated for 20 min at room temperature with gentle shaking. Biotinylated cells were then washed with PBS, resuspended in 250 ml of pre-warmed YPD medium and allowed to recover for 2 h at 30°C with shaking. Estradiol was added to a final concentration of 1 μM to induce the MEP (aging starts here). After 2 h of incubation at 30°C with shaking, 100 ml were harvested (young cells), while the rest of the culture (150 ml) was inoculated in a total volume of 1 L YPD containing 1 μM estradiol and 100 μg/ml ampicillin (to discourage bacterial contamination), and incubated at 30°C with shaking. Young cells were washed with cold PBS, resuspended in 5 ml cold PBS and incubated with 100 μl streptavidin-coated BioMag beads (Qiagen) in a 5 ml LoBind tube (Eppendorf) at 4°C with gentle shaking for 30 min. Cells were gently pelleted at 4°C (3 min 1800 × g), resuspended in 7 ml cold YPD and transferred to a glass test tube (Lab Logistics Group). The tube was placed in a magnet (“The Big Easy” EasySep Magnet, Stemcell Technologies) for 5 min on ice. Cells were then washed three times by removing supernatant by pipetting, resuspending them in 7 ml cold YPD and incubating for 5 min on ice in the magnet. Finally, cells were resuspended in 5.2 ml PBS and transferred in a 5 ml LoBind tube (Eppendorf). Of the 5.2 ml of purified mother cells, 100 μl were stained for bud scars counting (see below); 100 μl were diluted 1000x and plated on SD medium to assess cell viability; the remaining 5 ml were pelleted and plated on canavanine-containing SD medium (50 μg/ml) to identify forward mutations in *CAN1*. After 20 h of MEP induction, the entire aged 1 L culture was harvested. The aged cells were processed similarly to the young cells, with slight modifications because of the higher number of cells due to the presence of daughter cells. For beading, the cells were split into 4 different 5 ml LoBind tubes, and 50 μl streptavidin coated BioMag beads were added to each tube. For magnetic sorting, two glass tubes were used and cells were washed four times.

For both young and aged samples, colonies were counted after 2 d of growth at 30°C and the spontaneous forward mutation frequencies at the *CAN1* locus were determined. Expected mutation frequencies in aged cells were calculated as previously described [29].

### Bud scar detection and counting

Purified mother cells (see above) were stained with propidium iodide (PI) (Sigma) to identify viable cells and with Calcofluor White (Fluorescent Brightener 28, Sigma) to detect bud scars. 100 μl of purified mother cells in PBS (∼5 x 10^5^ cells) were stained with 2 μl of a 2 mM PI (Sigma) solution for 30 min at 30°C. Cells were then washed with ddH_2_O, fixed in 500 μl of 3.7% formaldehyde for 30 min at room temperature, washed with PBS, resuspended in 100 μl PBS and stored at 4°C. Just before imaging, cells were stained with Calcofluor White for 5 min at room temperature, washed with PBS and resuspended in 5-10 μl PBS. Images were acquired using a DeltaVision Elite imaging system (Applied Precision (GE), Issaquah, WA, USA) composed of an inverted microscope (IX-71; Olympus) equipped with a Plan Apo 100X oil immersion objective with 1.4 NA, InsightSSITM Solid State Illumination, excitation and emission filters for DAPI and A594, ultimate focus and a CoolSNAP HQ2 camera (Photometrics, Tucson, AZ, USA). Stacks of 30 images with 0.2 μm spacing were taken at an exposure time of 5 ms at 10% intensity for DAPI (Calcofluor White staining) and 50 ms at 32% intensity for A594 (PI staining). Reference bright-field images were also taken. Fluorescent images were subjected to 3D deconvolution using SoftWoRx 5.5 software (Applied Precision). Processing of all images was performed using Fiji (ImageJ, National Institute of Health) [98]. Bud scars from at least 50 PI-negative cells (which were alive after magnetic sorting) were manually counted for each sample to determine the cells’ replicative age.

### Gene Ontology enrichment analysis and functional annotation

GO enrichment analysis was performed with DAVID 6.8 (https://david.ncifcrf.gov/home.jsp) using the Functional Annotation tool [104,105]. To reduce functional redundancy among GO terms, we used the REVIGO Web server (http://revigo.irb.hr/) with a cutoff value C = 0.5 [106].

Functional enrichment within the yeast global genetic similarity network was performed and visualized with TheCellMap.org (http://thecellmap.org/), using SAFE [107,108].

## Acknowledgments

We thank Marlien Visser for helping with the fluctuation tests and Grant W. Brown for critically reading the manuscript.

## Supporting information

**S1 Fig. Exclusion of strains that escaped before the beginning of the screen.** When serial dilutions of strains from the MEP-YKO collection are spotted in the presence of estradiol, growth of MEP-proficient strains is restricted, while escaper strains grow normally. An example of one MEP-proficient strain and one escaper is shown.

**S2 Fig. Raw data for all the age-dependent mutation frequency measurement experiments.** (A) Mutation frequencies at the *CAN1* locus in young and old cells from the indicated strains. In the case of old cells, both observed and expected mutation frequencies are shown. For each individual experiment, the median replicative age of the young and old cell populations is indicated. (B) Bud scars distribution of young and old cell populations from each experiment.

**S3 Fig. *ICE2, ATG23* and *ROX3* did not validate as age-specific mutation suppression genes.** (A) *CAN1* forward mutation rate in young wt, *ice2Δ, atg23Δ*, and *rox3Δ* cells. Mean values from three independent experiments are plotted. Error bars represent standard error. ns: non-significant. (B) Age-dependent mutation frequencies at the *CAN1* locus in wt (replicative age ∼17), *ice2Δ* (replicative age ∼15), *atg23Δ* (replicative age ∼15.5), and *rox3Δ* (replicative age ∼15) cells. The difference between observed and expected mutation frequency is plotted. For the wt, the mean value from four independent experiments is plotted. Error bars represent standard error. For the mutants, only one experiment is shown (see the Results and Material and Methods sections for details).

**S4 Fig. *PEX3* deletion results in elevated spontaneous mutations in aged cells.** (A) *CAN1* forward mutation rate in young wt, and *pex3Δ* cells. Mean values from three or four independent experiments are plotted. Error bars represent standard error. ns: non-significant. (B) Age-dependent mutation frequencies at the *CAN1* locus in wt (replicative age ∼17), and *pex3Δ* (replicative age ∼13) cells. The difference between observed and expected mutation frequency is plotted. For the wt, the mean value from four independent experiments is plotted. Error bars represent standard error. For *pex3Δ*, only one experiment is shown.

**S1 Table. ScreenTroll phenotypic enrichment analysis.** ScreenTroll analysis (http://www.rothsteinlab.com/tools/screenTroll) identifies overlaps between published screens and our gene dataset (cutoff 75% escaper frequency, determined by CLIK). The top overlaps are related to genome instability and DNA damage sensitivity. “ORFs in screen” refers to the total number of hits identified in the overlapping screen.

**S2 Table. List of putative age-specific mutation suppression genes.**

**S3 Table. Escaper formation is not always caused by mutations at the *cre-EBD78* locus.**

**S1 File. Complete escaper frequency data from the screen and validation of mutation suppression genes.**

**S2 File. SAFE enrichments and GO enrichment analysis for general and MEP-specific mutation suppression genes.**

